# Forecasting drug resistant HIV protease evolution

**DOI:** 10.1101/2025.03.31.646462

**Authors:** Manu Aggarwal, Vipul Periwal

**Affiliations:** National Institutes of Health, Bethesda, MD

**Keywords:** HIV treatment, statistical physics, protein evolution, drug resistance

## Abstract

Protease inhibitors (PIs) target the protease (PR) enzyme to suppress viral replication. Their efficacy in human immunodeficiency virus treatment is compromised by the emergence of drug-resistant strains. Therefore, forecasting drug-resistance during viral evolution would help in the design of effective treatment strategies. We develop a probabilistic large-deviation model to infer epistatic interactions in genotypes observed in different treatment regimens and compute transition probabilities of point-mutations conditioned on the genotype and the treatment regimen. We simulate stochastic evolutionary paths weighted by such transition probabilities and show that low probable mutations are required for the viral population to evolve to diverse fit genotypes. We train classification models using a clinical data set of *in vitro* susceptibility tests to learn to infer the drug resistance of a genotype. We infer drug resistance along simulated evolutionary paths and predict that the combination PI-therapy of Atazanavir (ATV) and Ritonavir (RTV) is the least drug resistant. Without prior knowledge of PI-associated mutations, our model predicts known primary and secondary PI-resistant mutations as critical to drug resistance. This validates that our model learned mechanistic relations in the small data sets, tackling the challenge of sparse sequence data compared to the large combinatorial complexity of protein evolution and changing functionality in dynamic environments.

## 1 Introduction

Human immunodeficiency virus (HIV) antiretroviral therapy (ART) uses a combination of antiretroviral drugs for ongoing suppression of viral replication. Protease inhibitors (PIs) are a crucial component of ART that target the essential HIV protease (PR) enzyme. However, HIV’s high replication rate (10^7^–10^9^ newly infected cells/day in a patient [Coffin and Swanstrom, 2013]) facilitates the emergence of drug-resistant strains under the selective pressures of PI therapies that compromise their efficacy. For example, drug resistance to protease inhibitors (PIs) has been shown to emerge in up to 50% of patients [Richman et al., 2004]. Therefore, it would be helpful to forecast viral evolutionary paths that lead to drug-resistant strains in different ART treatment regimens. However, forecasting protein evolution faces the following challenges.

First, we do not have adequate mechanistic knowledge of the processes that drive evolution. A viral evolutionary path is a sequence of evolving protein genotypes that is directed by the deterministic force of natural selection and the stochastic influences of mutations and genetic drift [Lenski et al., 1991, Kryazhimskiy et al., 2014]. There is no consensus on the mechanism underlying natural selection because it is defined as a function of *fitness* of different genotypes [Orr, 2009] and there is no broad consensus on the precise biological and mathematical definition [Barker, 2009]. Previous studies have correlated viral fitness with different properties, for example, protein structural stability, replicative ability, epidemiological fitness, transmissive ability, and enzymatic activity.

Second, the effects of a mutation on the properties of a genotype depend on the selective pressures in the environment [Dolan et al., 2018] and also on the genotype or the sequence background in which the mutation is introduced [Starr and Thornton, 2016]. Due to the latter phenomenon, called epistasis, a specific point-mutation can theoretically have up to 20^*L*^ different effects on protein genotypes of length *L* since there are 20 different amino acids. Therefore, even in the same environment or set of selective pressures of a specific ART treatment regimen, a large number of evolutionary paths are possible due to the combined phenomena of stochastic influences and epistasis.

Our first contribution is a probabilistic large-deviation model that addresses these challenges in forecasting evolution. Our model simulates evolutionary paths in different treatment regimens using the free energy minimization (FEM) method [Hoang et al., 2019], a statistical physics based method of model inference that learns mechanistic relations in data. Crucially, FEM infers potentially asymmetric epistatic interactions as it does not make assumptions about statistical equilibrium, and can therefore be used to learn the coevolutionary information in a set of related protein sequences and infer transition probabilities of point-mutations conditioned on the genotype. Coevolutionary information has previously been related to protein structure, function, and fitness [De Juan et al., 2013, Stein et al., 2015, Serohijos and Shakhnovich, 2014, Neuwald, 2016]. Our approach avoids a mechanistic formulation of natural selection and fitness. It takes the stochasticity of evolution into account: mutations are randomly selected but weighted by inferred transition probabilities. We take into account the selective pressures of different PI treatment regimens by training FEM on sets of related protein sequences that are isolated from subjects who received the same set of PIs.

Probabilistic models based on statistical physics have previously been shown to perform well in extracting coevolutionary information from finite observed samples of related protein sequences. For example, Ferguson et al. [2013] and Levy et al. [2017] used the infinite-range Ising spin glass model [Binder and Young, 1986] and the Hamiltonian Potts model [Mora and Bialek, 2011], respectively, to predict the probabilities of observing specific mutation patterns that were consistent with the observed sequence statistics. FEM fundamentally differs from these approaches as follows. First, sets of related protein sequences from different treatment regimens that were available to us have a very small number of samples, a few hundred in most cases. Crucially, FEM has been shown to perform well for small data sets, for example, up to four times lower inference error compared to maximum likelihood estimation [Hoang et al., 2019]. Second, we directly infer epistatic interactions by computing the conditional probability of observing a particular residue at a position in a given genotype. Third, we do not use an energy model because that assumes an equilibrium distribution. This is critical because large deviations in the genotype are crucial in viral evolution to adapt to changing selective pressures. Hence, it cannot be expected that there is a static equilibrium distribution of viral genotypes during evolution. Consequently, the non-equilibrium approach of FEM does not assume symmetry and infers asymmetric epistatic relations—the effect of position *i* on *j* is not the same as the effect of *j* on *i*. Fourth, unlike previous approaches, we simulate evolutionary paths as we infer transition probabilities conditioned on the genotype. This allows us to compute multiple characteristics of evolutionary paths in different ART treatment regimens, for example the importance of rare events for genetic drift and the probability to reach drug resistance.

The second contribution of this work is a framework that couples two data sets from the publicly available Stanford HIVDB [Rhee et al., 2003, Shafer, 2006] to forecast the emergence of drug-resistant PR strains that have high FEM-fitness, defined specifically and precisely for our approach in the following, and high probability of emergence under different PI treatment regimens. It also determines mutations critical to the emergence of such strains. The first data set is the genotype-rx correlation data set that lists PR isolates from subjects and the PIs they received. We train FEM-based models to infer conditional point-mutation probabilities in the different treatment regimens and simulate evolutionary paths. We define FEM-fitness of a genotype as its likelihood to mutate, computed using the inferred point-mutation probabilities. The second data set is the genotype-phenotype correlation data set that lists PR isolates with their clinically measured drug fold resistance to different PIs. We used this data set to train FEM-based classification models to learn to infer drug resistance of PR sequences along the simulated evolutionary paths. Finally, we show that prohibiting certain mutations significantly reduces the chances of emergence of fit drug-resistant strains.

### 2 Results

Given a PR sequence or genotype ***σ*** = (*σ*_1_, …, *σ*_*L*_), we infer the probability of the amino acid at position *i*, or *A*_*i*_, to be *α* given the amino acids at all other positions in ***σ*** when treated with one or more PIs from Atazanavir (ATV), Ritonavir (RTV), Indinavir (IDV), Lopinavir (LPV), Nelfinavir (NFV), and Saquinavir (SQV). We denote this probability by *P*_*F*_ (*A*_*i*_ = *α* | ***σ***_∼*i*_) where *F* is the set of PIs or treatment regimen received by the subject. We define *F* = {None} for no treatment or no PIs received. We define the log likelihood of *σ* to mutate in the treatment regimen *F* as *m*_*F*_ (***σ***) = − min_1 ≤ *i* ≤ *L*_ {log(*P*_*F*_ (*A*_*i*_ = *σ*_*i*_ | ***σ***_∼*i*_))}.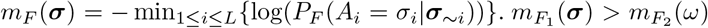 implies that the sequence ***σ*** is more likely to mutate in the treatment regimen *F*_1_ than *ω* in the treatment regimen *F*_2_, and we say that ***σ*** has a lower FEM-fitness in *F*_1_ than *ω* in *F*_2_.

### 2.1 PR is more likely to mutate under PI treatments

We visualize PR sequences as points in a two-dimensional space defined by the first two components of the transformations of their onehot-encodings using principal component analysis (PCA). Figure 1A shows the resulting 2D embedding of all PR sequences belonging to different treatment regimens. We call a PR sequence drug-naive if it belongs to no treatment. Figure 1B shows the embedding of the drug-naive sequences. Figure 1C shows an interpolated heatmap of the FEM-fitness, or FEM-fitness landscape, of the drug-naive sequences when there is no PI in the environment. Figure 1D shows the FEM-fitness landscapes of the drug-naive sequences in different PI treatments. Kernel density estimation (KDE) plots in Figure 1E show that drug-naive PR sequences are more likely to mutate when subjects are treated with PIs.

**Figure 1:**
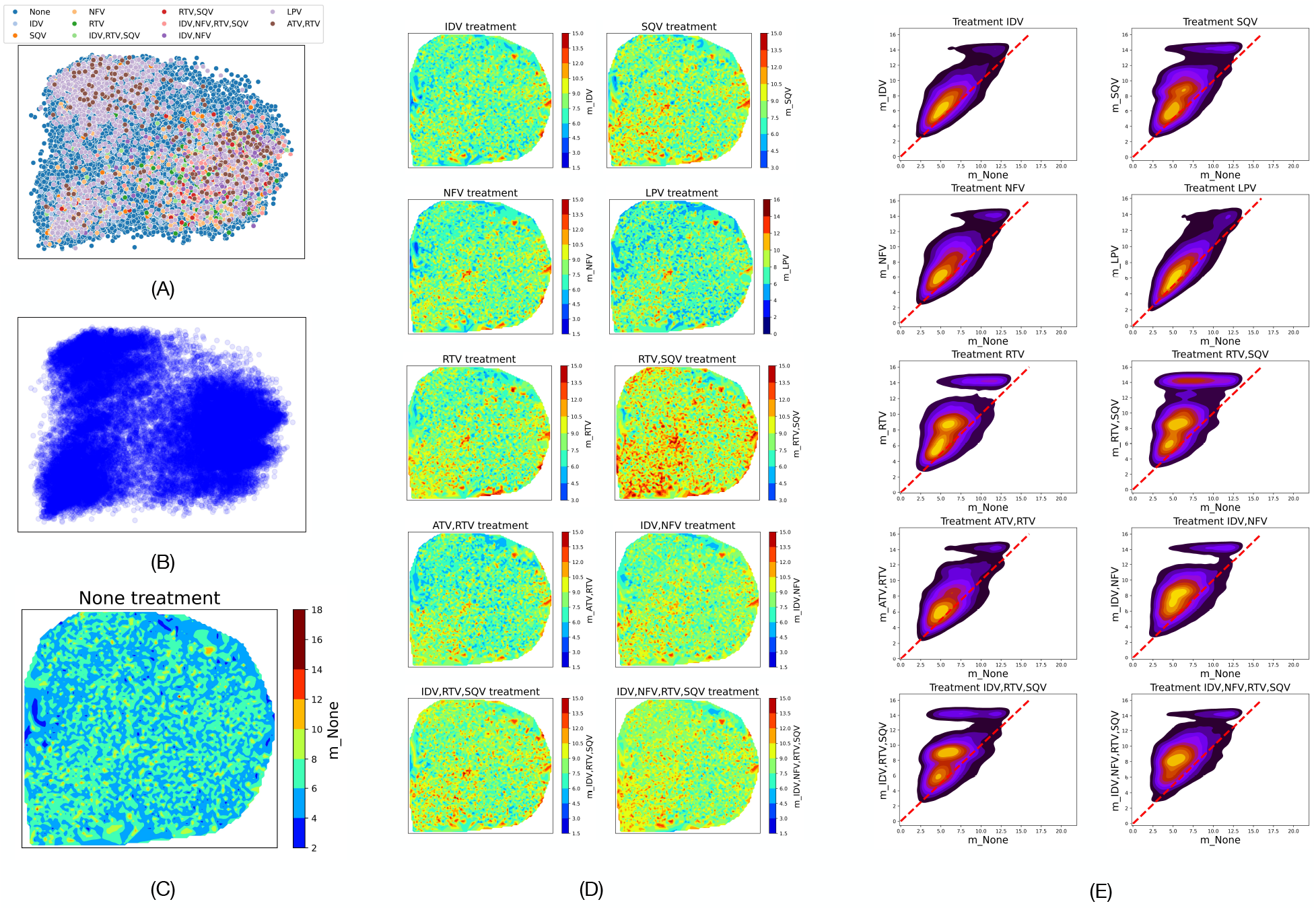
PR is more likely to mutate under PI treatments. (A) 2D embedding of PR isolates from subjects, colored by the treatment regimen they received. (B) 2D embedding of drug-naive isolates. (C) Negative log likelihoods of the drug-naive genotypes to not mutate when no PIs are given. (D) Negative log likelihoods of the drug-naive genotypes to not mutate in different treatment regimens with at least one PI. (E) Comparing negative log likelihood of the drug-naive genotypes to not mutate when there is no PI(*x*-axis) vs. when at least one PI is given. 2D KDE shows that majority of genotypes have higher likelihood to mutate under PI therapy (they are above the *y* = *x* red dashed line).

### 2.2 Rare mutations enable PR evolution to explore diverse fit genotypes

We simulate evolutionary paths using conditional point-mutation probabilities and use importance sampling to observe effects of rare mutations without having to simulate an excessive number of paths. Briefly, given an initial protein sequence ***σ*** = (*σ*_1_, …, *σ*_*L*_), we stochastically select a point-mutation weighted by 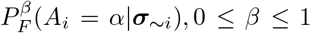, to result in a new sequence, say 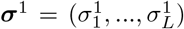. *β* is the hyperparameter for importance sampling that we call the inverse temperature. A lower value of *β* will increase the chances of selecting rare mutations. We compute the conditional probabilities for the new sequence and repeat the process iteratively *t* times, resulting in a path of protein evolution *T*^*F*^ = (***σ, σ***^1^, …, ***σ***^*t*^). The starting sequence for all simulations is the PR consensus subtype B sequence. We visualize the evolutionary paths in a 2D space by embedding the sequences in the paths using the first two components of the PCA transformations of their onehot-encodings.

Figure 2A shows the distributions of the probabilities of mutations selected at different values of *β*. A higher number of less probable mutations are selected at lower values of *β*. Figure 2B shows that evolutionary paths spread more at lower values of *β* and explore a greater number of different genotypes. This may not be consequential if the new genotypes explored at lower *β* values have low FEM-fitness. However, analysis reveals a nontrivial pattern. Figure 2C shows the − log likelihood to not mutate of the genotypes explored by simulated evolutionary paths at different values of *β*. The counts of genotypes of high FEM-fitness (− log likelihood to not mutate *<* 5) is higher at higher values of *β* and those with low FEM-fitness (− log likelihood to not mutate *>* 10) is higher at lower values of *β*. This is expected since selecting highly probable mutations (high *β*) results in fit genotypes and selecting low probable mutations (low *β*) results in unfit genotypes. However, the relationship between the counts of genotypes of FEM-fitness between − 10 and − 5 is not a monotonic function of *β*—the number of these genotypes at *β* = 0.6 and 0.8 is higher by an order of magnitude compared to when *β* is 0.4 and 1.0. Therefore, the rare events at *β* = 0.6 and 0.8 enable the evolutionary paths to explore an exponentially higher number of sequences.

**Figure 2:**
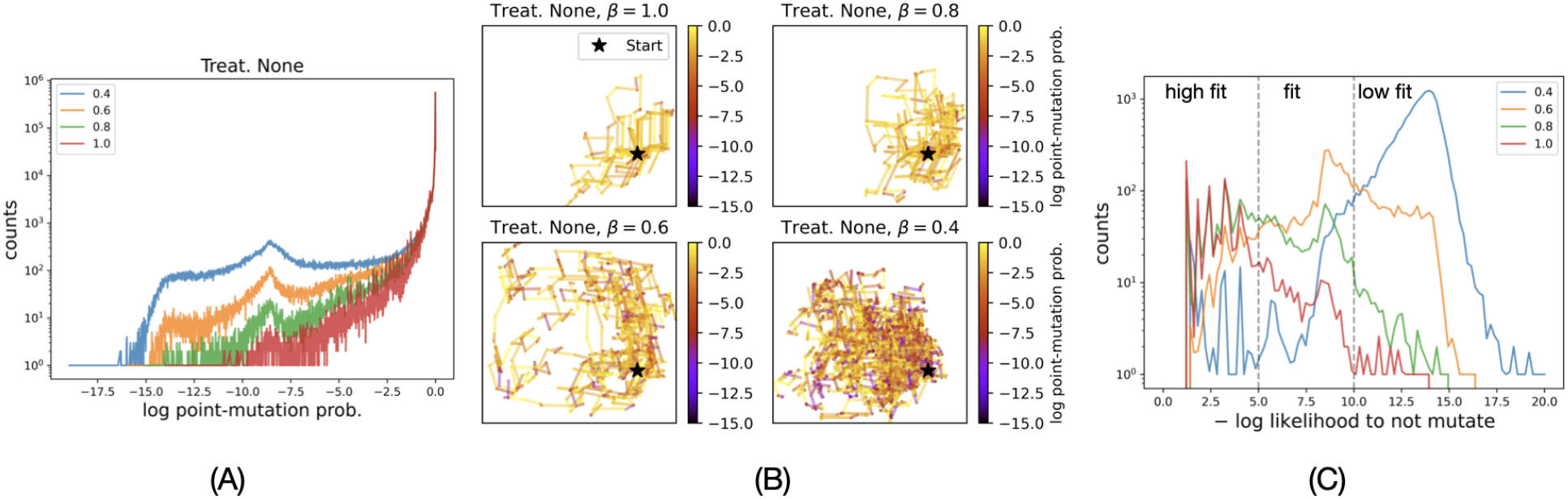
Low probable mutations enable PR evolution to explore diverse genotypes. (A) At lower *β* values, more number of mutations with log mutation probability less than *<* − 10 are explored by the simulated evolutionary trajectories. (B) 2D embeddings of 10 simulated trajectories, each of 1000 steps, at different *β* values when no PI is given. (C) Distribution of − log likelihoods to not mutate of unique genotypes explored by 1000 evolutionary paths simulated at different values of *β*. At *β* = 0.4 and 0.6, evolutionary paths explore more fit genotypes as compared to 0.2 and 1.0.

**Figure 3:**
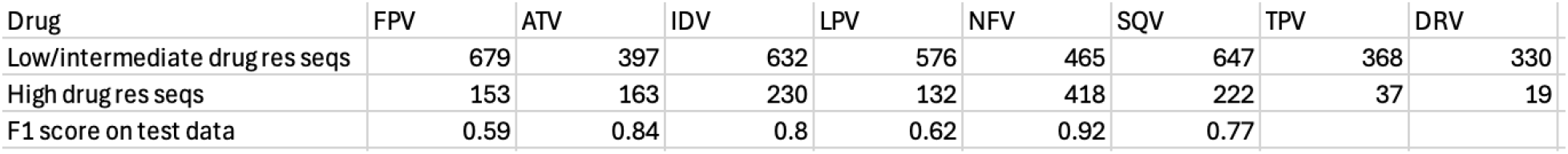
Number of unique genotypes in the classes of high and not-high drug resistance to 8 different drugs. FEM models were trained to infer drug resistance to drugs that had at least 100 samples in both classes of drug-resistance. F1-scores of the trained FEM models on test data are shown.

### 2.3 FEM inference of drug resistance achieves an accuracy F1-score of 0.92 on test data

We labeled high drug resistant genotypes in the genotype-phenotype data set with 1, and 0 otherwise (see Methods). We trained FEM models to infer high drug resistance of a PR sequence to 6 different PIs—Atazanavir (ATV), Fosamprenavir (FPV), Indinavir (IDV), Lopinavir (LPV), Nelfinavir (NFV), and Saquinavir (SQV). We ignored Tipranavir (TPV) and Darunavir (DRV) because the database had fewer than 100 isolates that showed high resistant to these PIs. Train/test split was 80 : 20. Table 3 shows the counts of different classes and the F1-scores of the trained FEM models on the test set. Even with fewer than a thousand samples and imbalanced data sets, FEM achieves high F1-scores of more than 0.8 for inference of high resistance to NFV, IDV, and ATV. Subsequently, we infer the resistance of PR genotypes to these three drugs in simulated evolutionary paths under different treatment regimens.

### 2.4 Treatments with multiple PIs have higher chances of emergence of highly fit drug resistant PR strains

We infer drug resistance of the genotypes in simulated evolutionary paths using FEM models trained on the genotype-phenotype data set. Figure 4A shows high NFV-resistant genotypes (marked red ×) in the evolutionary paths simulated under the treatment regimen *F* = {NFV} for different values of *β*. For each NFV-resistant genotype ***σ***, we compute − *m*_*F*_ (***σ***) and the maximal probability of reaching it along the simulated evolutionary paths in the treatment regimen *F* (see Methods). Figure 4B plots these two properties of all NFV-resistant sequences in the simulated evolutionary paths for all four values of *β* in the treatment regimen {NFV}. Figure 4C shows a 2D histogram of the same.

**Figure 4:**
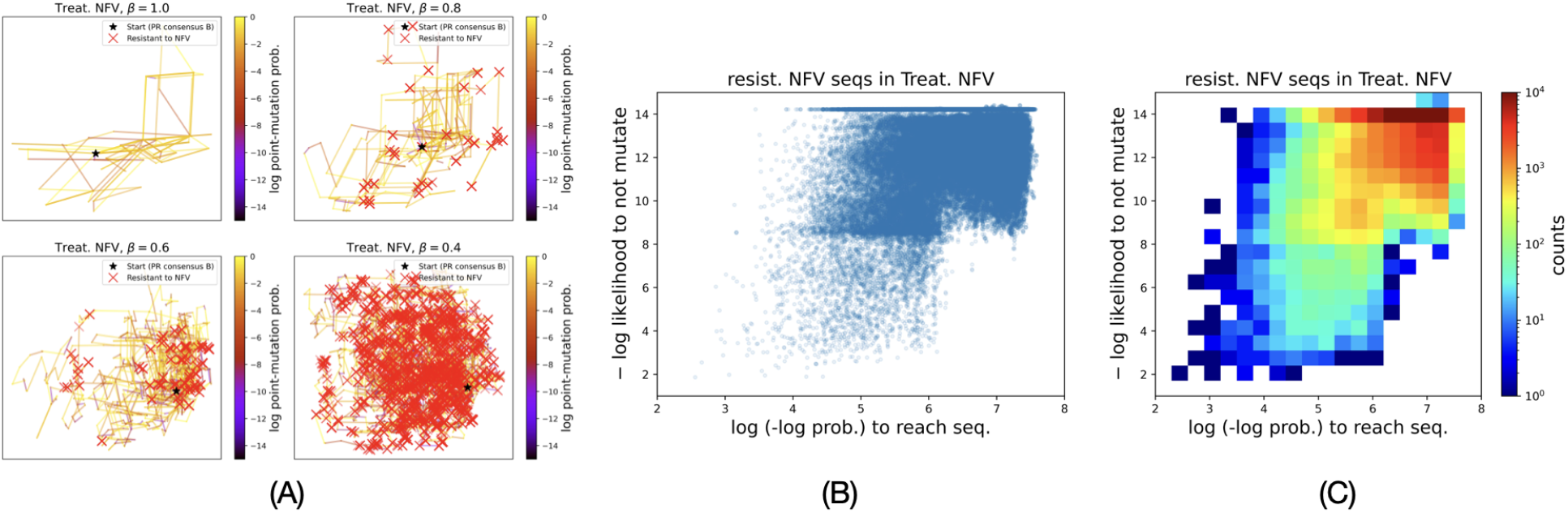
Coupling protein evolution and drug-resistance. (A) 10 simulated evolutionary paths at different values of *β* under the treatment regimen of {NFV} and red × mark the genotypes that were inferred to have high resistance to NFV. (B) Distribution of all drug resistant genotypes explored by simulated evolutionary paths for all *β* values combined. 1000 paths of 1000 steps were simulated for each *β* value. (C) 2D histogram of the distribution of drug resistant paths to show counts.

We analyze the effects of treatment regimens on the FEM-fitness of drug resistance strains as well as the likelihood of their emergence during evolution. We use Pareto optimality to determine strains that have optimal values of both criteria (see Methods). Briefly, we say that a genotype has Pareto optimality *ϕ* if the L_2_ norm of the log likelihood of not mutating and the log log probability of reaching is *ϕ*. The features are rescaled to [0, 1] by dividing them by respective upper bounds of 20 and 10. Hence 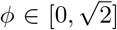. Figure 5A shows drug resistant genotypes (marked red) at different thresholds for *ϕ*. A lower *ϕ* (top left panel) implies drug resistant sequences with higher FEM-fitness and higher probability of reaching by the simulated evolutionary paths. Figure 5B plots the cumulative counts of drug-resistant genotypes as a function of thresholds for *ϕ*.

**Figure 5:**
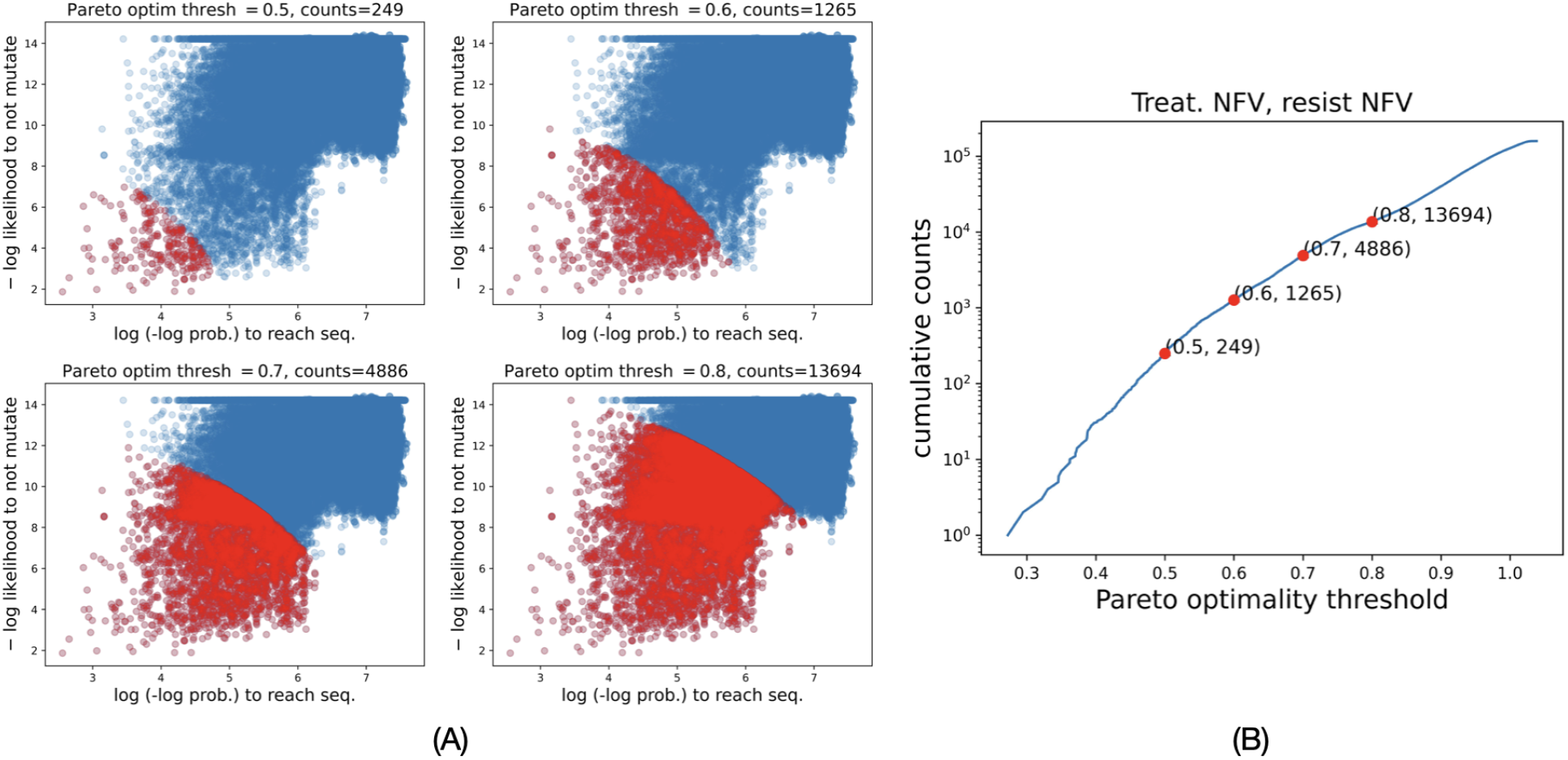
Pareto optimal measure of FEM-fitness of a drug resistant genotype. (A) Minimizing both − log likelihood to not mutate and log(− log prob. to reach) defines genotypes with higher FEM-fitness and higher chances of being explored by evolutionary paths. Pareto optimality threshold (*ϕ*) is the threshold on the norm of the two criteria. Red points mark genotypes within different Pareto optimality thresholds. (B) Cumulative counts of genotypes as a function of *ϕ*.

Figures 6A-C top panels show the cumulative counts of drug-resistant genotypes as functions of thresholds for *ϕ* for the different treatment regimens. The black-dashed line shows the baseline case when there was no treatment. We compare the fractional change in the counts for different treatment regimens with respect to the no-treatment regimen in Figures 6A-C bottom panels. PI treatments increase the number of drug resistant sequences by multiple orders of magnitude, more so for 0.4 *< ϕ <* 0.8. Hence, the number of sequences at thresholds lower than the maximal value of *ϕ* increases disproportionately, indicating that PI treatments result in a larger number of drug-resistant genotypes that have a higher FEM-fitness and the simulated evolutionary paths had a higher probability of reaching. Specifically, treatment regimens with a larger number of different PIs result in a higher chance of emergence of fit drug-resistant strains.

**Figure 6:**
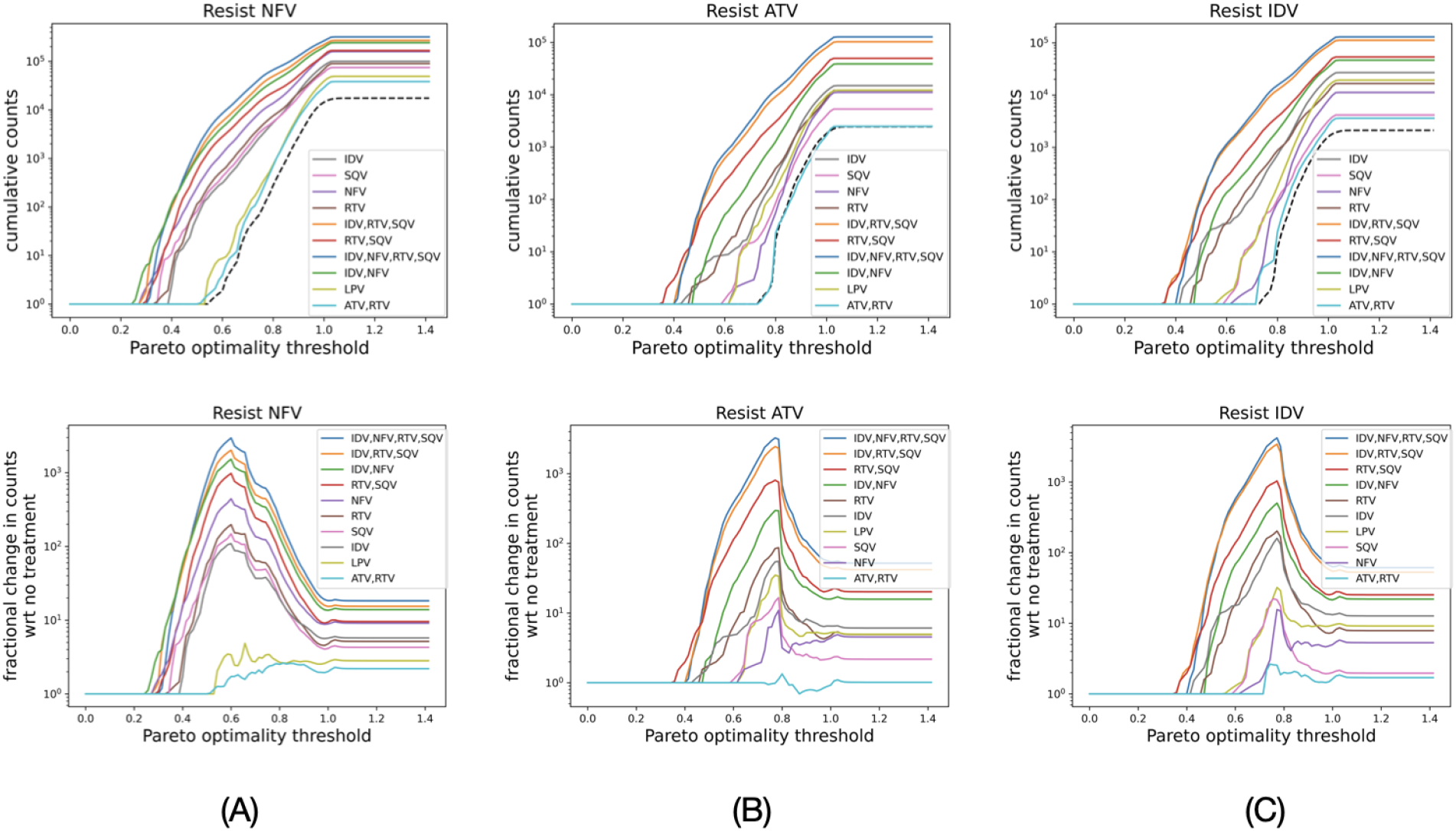
PI therapy significantly increases percentage of fit drug-resistant strains that have high chances to emerge during evolution. (top panel) Cumulative counts of drug resistant sequences as a function of *ϕ*. Black dashed plot shows no-treatment or when no PIs were given. (bottom panel) Fractional change in the cumulative counts with respect to when no PI is given. The legends in the bottom panel are in the decreasing order of the peaks of the curves.

### 2.5 The FEM framework infers mutations critical to the emergence of fit drug resistant strains

We determine the importance of a point-mutation in the emergence of drug resistance by analyzing the fractional change in the counts of drug resistant genotypes as a function of *ϕ* when a point-mutation is not allowed with respect to when all mutations are allowed. Figure 7 shows results for treatment regimens with multiple PIs. We do not show {ATV, RTV} treatment in this figure because the fractional change in the number of fit drug-resistance sequences with respect to no-treatment is not significant (Figure 6 (lower row) {ATV, RTV} plots). We identify and label outlier curves in the figures on the basis of visual comparison. In most cases, prohibiting these point-mutations reduces the number of drug resistant genotypes by more than 90% (fraction value less than 0.1). Importantly, this significant reduction is in the region 0.3 *< ϕ <* 0.6 (vertical dashed lines in the figure), which is lower than the maximum value of 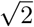 of *ϕ*. Hence, these point-mutations are critical to the emergence of drug-resistant strains that have high FEM-fitness.

**Figure 7:**
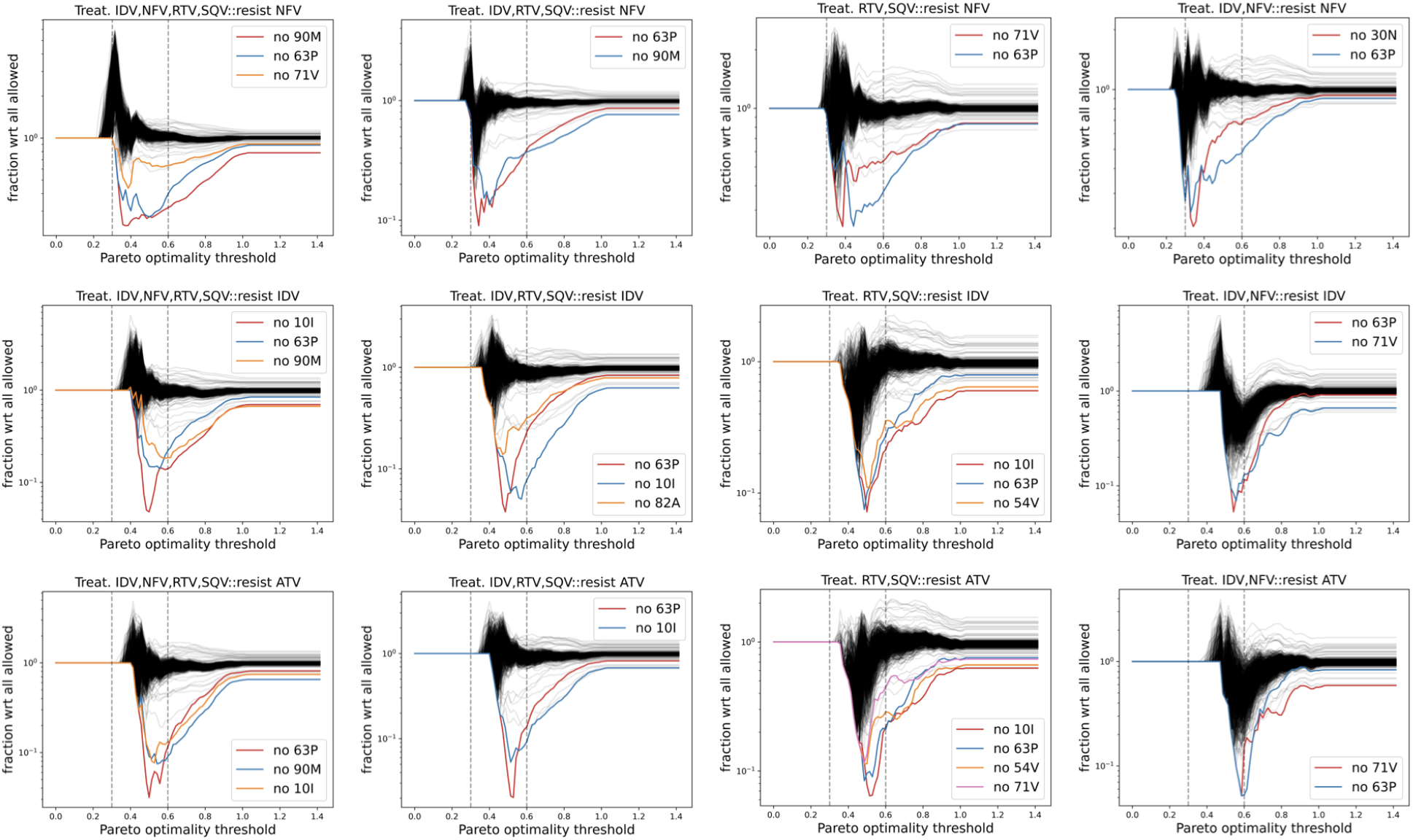
Mutations 90M, 10I, 30N, 63P, 54V, 71V, and 82A are critical to higher chances of emergence of fit drug-resistant strains. Prohibiting any of these mutations significantly reduces the percentage of genotypes (in most cases more than 90% reduction) in the region of 0.3 *< ϕ <* 0.6 (vertical dashed lines). This region corresponds to relatively fit genotypes since the maximum value of *ϕ* is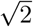.

## 3 Discussion

We introduced a probabilistic model to directly infer potentially asymmetric epistatic interactions and transition probabilities in a given sequence background and under different PI treatment regimens. We defined the FEM-fitness of a PR genotype as a function of the genotype and the selective pressure of the combination of PIs received by the subject from which it was isolated. We simulated evolutionary paths and showed that low probable mutations or rare events significantly increase the chances of evolving to fit genotypes. We then used FEM to infer the drug resistance of PR genotypes to different PIs. We combined simulated evolutionary paths with drug-resistance inference to reveal that the chances of emergence of fit drug-resistant genotypes increase with the number of PIs in the treatment regimen, with the exception of {ATV, RTV}. Finally, we analyze the effects of prohibiting mutations and determine 90M, 10I, 30N, 63P, 54V, 71V, and 82A as critical to higher chances of emergence of drug-resistant strains of high FEM-fitness.

Mutations D30N, V82A, and L90M are among the primary causes of resistance to PIs as they arise within the active site of the enzyme, directly affecting inhibitor binding [Ali et al., 2010, Gulnik et al., 1995]. On the other hand, mutations I54V, L10I, and A71V are known as secondary mutations that can modulate the effect of primary mutations on phenotypic resistance [Servais et al., 2002]. Interestingly, 63P was determined as critical in all the cases shown in Figure 7 even though it is neither a primary nor a secondary drug-resistant mutation. It is a polymorphic mutation—a variation in the DNA sequence that naturally occurs (without treatment) in a population with a frequency of 1 % or higher [Brookes, 1999]. However, Bossi et al. [1999] reported 63P as the most common amino acid change in PR, observed in 44% of HIV-1 infected patients treated with PIs.

The publicly available data sets used in this work are limited to treatment regimens with single or multiple drugs from one of four drug classes—protease inhibitors (PIs), non-nucleoside reverse transcriptase inhibitors (NNRTIs), nucleoside reverse transcriptase inhibitors (NRTIs), and integrase strand transfer inhibitors (INIs). However, regimens consisting of three drugs of different classes, called HAART (Highly Active Antiretroviral Therapy), have been found to be more effective [Delaney, 2006, Vlahov et al., 2005, Yeni, 2006], with one of the advantages being a reduction in the emergence of drug resistance. Our approach could be applied to data from HAART treatment regimens to determine the mechanisms that alleviated drug resistance in this combination therapy, insights that may be useful in improving treatment regimens.

Recently, treatment regimens consisting of two drugs have been recommended due to their reduced adverse effects and toxicities compared to three-drug regimens [Gibas et al., 2022]. Our analysis predicts that the {ATV, RTV} treatment regimen has the lowest chances of emergence of drug resistance, even compared to mono-PI therapies (see Figure 6 bottom panel). However, as with all data-driven inference in complex biological processes, only clinical data can confirm the efficacy of this regimen compared with others.

## 4 Methods

### 4.1 Data

All data is taken from the publicly available Stanford HIV database [Rhee et al., 2003, Shafer, 2006]. The consensus sub-type B protease (PR) sequence is taken from https://hivdb.stanford.edu/pages/documentPage/consensus_amino_acid_sequences.html.

#### 4.1.1 Genotype-rx data set

Data set of protease isolates from different subjects (https://hivdb.stanford.edu/download/GenoRxDatasets/PR.txt). The treatment given to each subject prior to PR isolation is reported for each isolate as a set of protease inhibitors (PIs), which is a subset of {Atazanavir (ATV), Ritonavir (RTV), Indinavir (IDV), Lopinavir (LPV), Nelfinavir (NFV), Saquinavir (SQV)}. If no PI was given, it is reported as ‘None’. We processed the data set by removing PR sequences in three cases. (1) Isolates with insertions, deletions, or gaps in their sequence. (2) Isolates with ambiguity in their sequence—a mixture of amino acids reported at a position in the sequence. (3) Isolates with ambiguity in the treatment received—’Unknown’ or ‘PI’ in the reported treatment. Supplemental table shows treatment regimens with at least 100 unique PR isolates.

#### 4.1.2 Genotype-phenotype drug resistance data set

We used the genotype-phenotype correlation data set of PR isolates available at https://hivdb.stanford.edu/pages/genopheno.dataset.html. It reports *in vitro* drug-fold resistances [Zhang et al., 2005] of PR sequences to up to 8 different PIs—Atazanavir (ATV), Darunavir (DRV), Fosamprenavir (FPV), Indinavir (IDV), Lopinavir (LPV), Nelfinavir (NFV), Saquinavir (SQV), and Tipranavir (TPV). We ignored PR isolates with mixtures or X in any position to avoid ambiguous sequences. We annotated the resistance of a PR sequence to a drug as low/intermediate (label 0) or high (label 1) based on the high cut-off thresholds in Rhee et al. [2010]. We confirmed that there does not exist a PR sequence that was labeled both 0 and 1 with respect to the same drug.

### 4.2 Mathematical model

#### 4.2.1 Terminology

Let ℱ be the collection of treatment regimens in the processed genotype-rx data set and *S*_*F*_ be the set of PR sequences isolated from subjects given the treatment regimen *F* ∈ ℱ with |*S*_*F*_ | = *n*_*F*_,

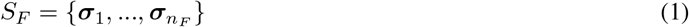

where

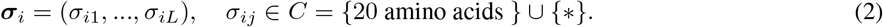

*L* = 99 for PR sequences.

Similarly, let *D* ={ATV, DRV, FPV, IDV, LPV, NFV, SQV, and TPV} and *S*_*d*_ be the set of PR sequences for which drug-fold resistance to *d* ∈ *D* is reported in the genotype-phenotype data set.

#### 4.2.2 Sequence onehot encoding

The set of all PR sequences in the data sets is *S* = (∪_*F*∈*ℱ*_ *S*_*F*_) ∪ (∪_*d*∈*D*_ *S*_*d*_). We denote the set of symbols at position *j* in the sequences in *S* by

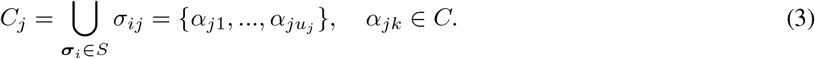

Supplemental table shows *C*_*j*_ for all positions.

Given a protein sequence ***σ*** = (*σ*_1_, …, *σ*_*L*_), the onehot encoding of *σ*_*j*_ is

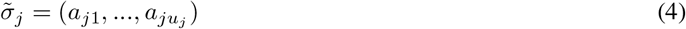

where

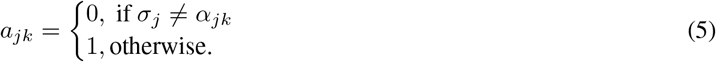

We denote onehot encoding of ***σ*** by

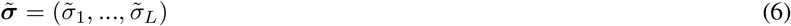

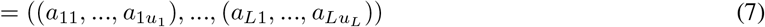

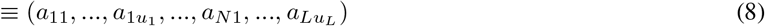

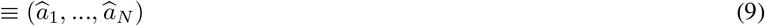

where *N* is the total number of bits in the onehot encoding,

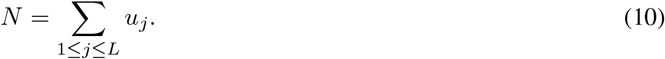

*N* = 1056 in this case.

#### 4.2.3 Free energy minimization (FEM)

Hoang et al. [2019] developed free-energy minimization (FEM), a statistical physics based method of model inference that learns mechanistic relations in data. Given a data set of *N* -dimensional binary observations each labeled 0 or 1, FEM computes a weight vector **w** = (*w*_1_, …, *w*_*N*_) ∈ ℝ^*N*^ and a bias *b* ∈ ℝ, and infers the probability that a *N* -dimensional binary sequence ***σ*** will be labeled *ω* ∈ {0, 1} as

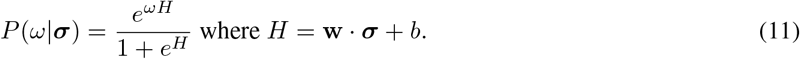

#### 4.2.4 transition probabilities of point-mutations conditioned on the genotype and treatment regimen

We denote sequence of amino acids at all positions except *p* by

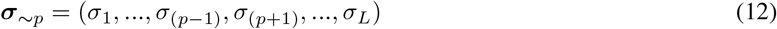

and its onehot encoding by

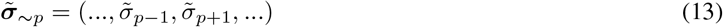

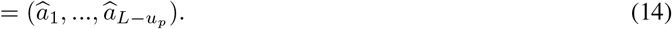

We denote the conditional probability of amino acid at position *p*, or *A*_*p*_, to be *λ* ∈ *C*_*p*_ given amino acids at all other positions and in the treatment regimen *F* by

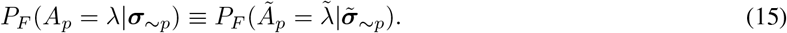

where

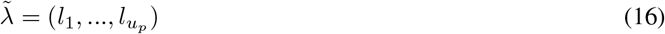

is the onehot encoding of *λ* at position *p*. We have

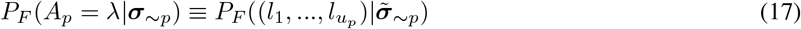

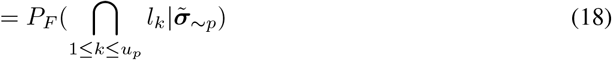

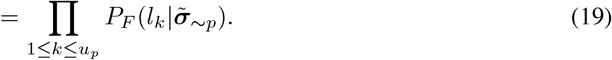

Using equation 11,

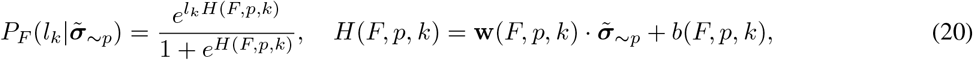

where the weight vector **w**(*F, p, k*) and the bias *b*(*F, p, k*) are computed by training FEM on the data set with inputs as onehot encodings 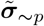 of sequences observed in treatment regimen *F* and with the corresponding output labels as the *k*-th bit of 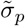.

#### 4.2.5 Inferring drug resistance

Resistance of a genotype ***σ*** to drug *d* is inferred as high if *P*_*d*_(1|***σ***) *>* 0.5 where

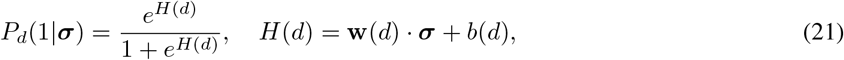

and weight vector **w**(*d*) and the bias *b*(*d*) are computed by training FEM on the data set with inputs as onehot encodings of sequences for which resistance to the drug *d* was labeled 0 or 1 as mentioned in section 4.1.2.

## Acknowledgements

This research was supported by the Intramural Research Program of the NIH, the National Institute of Diabetes and Digestive and Kidney Diseases (NIDDK). This work utilized the computational resources of the NIH HPC Biowulf cluster (https://hpc.nih.gov).

## Notes

### Competing Interest Statement

The authors have declared no competing interest.

