## Supplemental tables for "Forecasting drug resistant HIV protease evolution"

Table 1

| Treatment regimen | #seqs |
|---|---|
| None | 40820 |
| IDV | 558 |
| SQV | 155 |
| NFV | 514 |
| RTV | 150 |
| IDV,RTV,SQV | 153 |
| RTV,SQV | 107 |
| IDV,NFV,RTV,SQV | 179 |
| IDV,NFV | 166 |
| LPV | 1379 |
| ATV,RTV | 185 |

| Position | Unique amino acids (AA) observed | #AA |
|---|---|---|
| 1 | ['A', 'F', 'H', 'L', 'P', 'Q', 'R', 'S', 'T'] | 9 |
| 2 | ['*', 'E', 'H', 'I', 'K', 'L', 'P', 'Q', 'R', 'S', 'T', 'V'] | 12 |
| 3 | ['F', 'I', 'K', 'L', 'M', 'N', 'Q', 'S', 'T', 'V'] | 10 |
| 4 | ['A', 'C', 'D', 'F', 'H', 'I', 'K', 'L', 'N', 'P', 'Q', 'S', 'T'] | 13 |
| 5 | ['A', 'F', 'H', 'I', 'L', 'P', 'Q', 'R', 'S', 'T', 'V'] | 11 |
| 6 | ['*', 'C', 'E', 'G', 'L', 'P', 'R', 'S', 'W'] | 9 |
| 7 | ['*', 'D', 'E', 'H', 'K', 'L', 'P', 'Q', 'R', 'S', 'T', 'Y'] | 12 |
| 8 | ['*', 'E', 'G', 'H', 'L', 'P', 'Q', 'R', 'W'] | 9 |
| 9 | ['A', 'G', 'H', 'L', 'P', 'Q', 'S', 'T'] | 8 |
| 10 | ['A', 'C', 'F', 'G', 'H', 'I', 'L', 'M', 'N', 'P', 'Q', 'R', 'S', 'T', 'V', 'Y'] | 16 |
| 11 | ['A', 'C', 'D', 'F', 'G', 'I', 'K', 'L', 'M', 'S', 'T', 'V'] | 12 |
| 12 | ['*', 'A', 'D', 'E', 'F', 'G', 'H', 'I', 'K', 'L', 'M', 'N', 'P', 'Q', 'R', 'S', 'T', | 20 |
| 13 | ['A', 'C', 'D', 'E', 'F', 'G', 'I', 'K', 'L', 'M', 'N', 'R', 'T', 'V', 'Y'] | 15 |
| 14 | ['A', 'D', 'E', 'G', 'H', 'I', 'K', 'L', 'M', 'N', 'Q', 'R', 'S', 'T', 'V', 'W', 'Y' | 17 |
| 15 | ['*', 'A', 'D', 'E', 'G', 'I', 'K', 'L', 'M', 'N', 'R', 'S', 'T', 'V', 'Y'] | 15 |
| 16 | ['A', 'C', 'D', 'E', 'G', 'K', 'N', 'Q', 'R', 'V', 'W', 'Y'] | 12 |
| 17 | ['A', 'C', 'D', 'E', 'G', 'K', 'N', 'R', 'S', 'T', 'V'] | 11 |
| 18 | ['*', 'A', 'E', 'G', 'H', 'I', 'K', 'L', 'M', 'N', 'P', 'Q', 'R', 'S', 'T', 'V', 'Y'] | 17 |
| 19 | ['A', 'E', 'F', 'G', 'H', 'I', 'K', 'L', 'M', 'N', 'P', 'Q', 'R', 'S', 'T', 'V', 'W' | 17 |
| 20 | ['*', 'C', 'E', 'G', 'I', 'K', 'L', 'M', 'N', 'Q', 'R', 'T', 'V', 'Y'] | 14 |
| 21 | ['A', 'D', 'E', 'G', 'K', 'N', 'Q', 'R', 'T', 'V'] | 10 |
| 22 | ['A', 'C', 'D', 'G', 'L', 'P', 'R', 'S', 'T', 'V'] | 10 |
| 23 | ['F', 'H', 'I', 'K', 'L', 'P', 'Q', 'S', 'V'] | 9 |
| 24 | ['*', 'F', 'I', 'L', 'M', 'P', 'S', 'V', 'W'] | 9 |
| 25 | ['D', 'E', 'F', 'G', 'H', 'K', 'N', 'V', 'Y'] | 9 |
| 26 | ['A', 'I', 'P', 'R', 'S', 'T'] | 6 |
| 27 | ['E', 'G', 'P', 'R'] | 4 |
| 28 | ['A', 'E', 'G', 'P', 'Q', 'T', 'V'] | 7 |
| 29 | ['A', 'D', 'E', 'G', 'H', 'N', 'V', 'Y'] | 8 |
| 30 | ['D', 'E', 'G', 'K', 'N', 'S', 'V', 'Y'] | 8 |
| 31 | ['A', 'G', 'I', 'P', 'R', 'S', 'T'] | 7 |
| 32 | ['A', 'E', 'F', 'I', 'L', 'M', 'T', 'V'] | 8 |
| 33 | ['F', 'I', 'K', 'L', 'M', 'S', 'T', 'V'] | 8 |
| 34 | ['*', 'A', 'D', 'E', 'G', 'H', 'K', 'L', 'N', 'Q', 'S', 'T', 'V'] | 13 |
| 35 | ['*', 'A', 'D', 'E', 'G', 'H', 'K', 'N', 'Q', 'R', 'S', 'T', 'V', 'Y'] | 14 |
| 36 | ['A', 'D', 'E', 'F', 'I', 'K', 'L', 'M', 'N', 'S', 'T', 'V'] | 12 |
| 37 | ['A', 'C', 'D', 'E', 'F', 'G', 'H', 'I', 'K', 'L', 'M', 'N', 'P', 'Q', 'R', 'S', 'T', | 20 |
| 38 | ['C', 'F', 'G', 'I', 'K', 'L', 'M', 'S', 'V', 'W'] | 10 |
| 39 | ['A', 'E', 'H', 'I', 'K', 'L', 'M', 'P', 'Q', 'R', 'S', 'T', 'V'] | 13 |

| | | |
|---|---|---|
| 40 | ['E', 'G', 'R', 'T'] | 4 |
| 41 | ['*', 'A', 'D', 'E', 'G', 'H', 'I', 'K', 'N', 'P', 'Q', 'R', 'S', 'T', 'V', 'Y'] | 16 |
| 42 | ['*', 'C', 'G', 'L', 'M', 'R', 'W', 'Y'] | 8 |
| 43 | ['*', 'A', 'E', 'G', 'I', 'K', 'M', 'N', 'Q', 'R', 'S', 'T', 'V'] | 13 |
| 44 | ['A', 'K', 'L', 'P', 'Q', 'R', 'S', 'T'] | 8 |
| 45 | ['*', 'I', 'K', 'L', 'M', 'N', 'P', 'Q', 'R', 'T', 'V'] | 11 |
| 46 | ['I', 'K', 'L', 'M', 'R', 'T', 'V'] | 7 |
| 47 | ['A', 'I', 'K', 'L', 'M', 'R', 'T', 'V'] | 8 |
| 48 | ['A', 'E', 'G', 'I', 'K', 'L', 'M', 'Q', 'R', 'T', 'V', 'W'] | 12 |
| 49 | ['A', 'E', 'G', 'K', 'R', 'V'] | 6 |
| 50 | ['F', 'I', 'L', 'M', 'N', 'S', 'V'] | 7 |
| 51 | ['A', 'E', 'G', 'K', 'R', 'V', 'W'] | 7 |
| 52 | ['A', 'D', 'E', 'F', 'G', 'R', 'S', 'V'] | 8 |
| 53 | ['C', 'F', 'I', 'L', 'S', 'V', 'W', 'Y'] | 8 |
| 54 | ['A', 'F', 'I', 'K', 'L', 'M', 'R', 'S', 'T', 'V'] | 10 |
| 55 | ['E', 'F', 'G', 'H', 'I', 'K', 'M', 'N', 'Q', 'R', 'T'] | 11 |
| 56 | ['A', 'G', 'I', 'K', 'L', 'R', 'T', 'V'] | 8 |
| 57 | ['*', 'G', 'I', 'K', 'N', 'Q', 'R', 'S', 'T'] | 9 |
| 58 | ['*', 'E', 'H', 'K', 'L', 'P', 'Q', 'R'] | 8 |
| 59 | ['C', 'D', 'F', 'H', 'M', 'N', 'Q', 'S', 'Y'] | 9 |
| 60 | ['A', 'D', 'E', 'G', 'H', 'K', 'N', 'R', 'S', 'T', 'Y'] | 11 |
| 61 | ['A', 'D', 'E', 'G', 'H', 'K', 'L', 'N', 'P', 'Q', 'R', 'S', 'Y'] | 13 |
| 62 | ['G', 'I', 'K', 'L', 'M', 'N', 'Q', 'R', 'T', 'V'] | 10 |
| 63 | ['A', 'C', 'D', 'E', 'F', 'G', 'H', 'I', 'K', 'L', 'M', 'N', 'P', 'Q', 'R', 'S', 'T', | 20 |
| 64 | ['F', 'I', 'K', 'L', 'M', 'P', 'R', 'T', 'V'] | 9 |
| 65 | ['A', 'D', 'E', 'G', 'H', 'I', 'K', 'N', 'Q', 'R', 'T', 'V', 'Y'] | 13 |
| 66 | ['C', 'E', 'F', 'I', 'L', 'M', 'N', 'S', 'T', 'V'] | 10 |
| 67 | ['*', 'A', 'C', 'D', 'E', 'F', 'G', 'H', 'K', 'L', 'M', 'N', 'Q', 'R', 'S', 'W', 'Y | 17 |
| 68 | ['D', 'E', 'G', 'K', 'Q', 'R', 'V', 'W'] | 8 |
| 69 | ['A', 'C', 'E', 'H', 'I', 'K', 'L', 'M', 'N', 'P', 'Q', 'R', 'S', 'T', 'Y'] | 15 |
| 70 | ['A', 'E', 'G', 'H', 'I', 'K', 'L', 'M', 'N', 'P', 'Q', 'R', 'S', 'T', 'V', 'Y'] | 16 |
| 71 | ['A', 'D', 'G', 'I', 'L', 'M', 'N', 'P', 'S', 'T', 'V'] | 11 |
| 72 | ['A', 'D', 'E', 'F', 'I', 'K', 'L', 'M', 'N', 'P', 'Q', 'R', 'S', 'T', 'V', 'W', 'Y'] | 17 |
| 73 | ['A', 'C', 'D', 'G', 'K', 'N', 'R', 'S', 'T', 'V'] | 10 |
| 74 | ['A', 'E', 'I', 'K', 'M', 'P', 'Q', 'R', 'S', 'T', 'V'] | 11 |
| 75 | ['A', 'G', 'I', 'L', 'M', 'V'] | 6 |
| 76 | ['E', 'F', 'I', 'L', 'S', 'V'] | 6 |
| 77 | ['A', 'E', 'G', 'I', 'K', 'L', 'M', 'Q', 'T', 'V'] | 10 |
| 78 | ['*', 'A', 'D', 'E', 'G', 'R', 'V', 'Y'] | 8 |
| 79 | ['A', 'D', 'E', 'H', 'L', 'N', 'P', 'Q', 'S', 'T'] | 10 |

| | | |
|---|---|---|
| 80 | ['A', 'H', 'I', 'P', 'Q', 'R', 'S', 'T', 'V'] | 9 |
| 81 | ['H', 'L', 'P', 'S', 'T'] | 5 |
| 82 | ['A', 'C', 'D', 'E', 'F', 'H', 'I', 'L', 'M', 'N', 'P', 'S', 'T', 'V'] | 14 |
| 83 | ['D', 'H', 'I', 'K', 'N', 'P', 'S', 'T', 'Y'] | 9 |
| 84 | ['A', 'C', 'I', 'K', 'L', 'M', 'T', 'V'] | 8 |
| 85 | ['F', 'I', 'L', 'M', 'N', 'T', 'V'] | 7 |
| 86 | ['*', 'C', 'E', 'G', 'K', 'R', 'V'] | 7 |
| 87 | ['*', 'E', 'G', 'I', 'K', 'P', 'Q', 'R', 'S', 'T', 'V'] | 11 |
| 88 | ['D', 'G', 'H', 'I', 'K', 'N', 'S', 'T', 'V', 'Y'] | 10 |
| 89 | ['A', 'C', 'E', 'F', 'I', 'K', 'L', 'M', 'P', 'R', 'T', 'V', 'W'] | 13 |
| 90 | ['*', 'C', 'F', 'L', 'M', 'S', 'V', 'W'] | 8 |
| 91 | ['A', 'C', 'G', 'I', 'K', 'L', 'N', 'P', 'S', 'T', 'V'] | 11 |
| 92 | ['*', 'A', 'E', 'G', 'H', 'K', 'L', 'M', 'N', 'P', 'Q', 'R', 'S', 'T'] | 14 |
| 93 | ['A', 'D', 'F', 'H', 'I', 'K', 'L', 'M', 'N', 'P', 'R', 'S', 'T', 'V', 'Y'] | 15 |
| 94 | ['A', 'C', 'D', 'E', 'G', 'H', 'N', 'Q', 'R', 'S', 'V', 'W'] | 12 |
| 95 | ['A', 'C', 'F', 'G', 'I', 'L', 'M', 'N', 'R', 'S', 'V', 'W', 'Y'] | 13 |
| 96 | ['A', 'D', 'I', 'L', 'N', 'P', 'S', 'T'] | 8 |
| 97 | ['*', 'F', 'I', 'K', 'L', 'P', 'Q', 'S', 'T', 'V'] | 10 |
| 98 | ['C', 'D', 'H', 'I', 'K', 'N', 'Q', 'R', 'S', 'T', 'Y'] | 11 |
| 99 | ['F', 'G', 'I', 'L', 'S', 'V', 'W', 'Y'] | 8 |
